# Landmark navigation in a mantis shrimp

**DOI:** 10.1101/2020.03.12.988741

**Authors:** Rickesh N. Patel, Thomas W. Cronin

## Abstract

Mantis shrimp are predatory crustaceans that commonly occupy burrows in shallow, tropical waters worldwide. Most of these animals inhabit structurally complex, benthic environments where many potential landmarks are available. Mantis shrimp of the species *Neogonodactylus oerstedii* return to their burrows between foraging excursions using path integration, a vector-based navigational strategy that is prone to accumulated error. Here we show that *N. oerstedii* can navigate using landmarks in parallel with their path integration system, offseting error generated when navigating using solely path integration. We also report that when the path integration and landmark navigation systems are placed in conflict, *N. oerstedii* will orient using either system or even switch systems enroute. How they make the decision to trust one navigational system over another is unclear. These findings add to our understanding of the refined navigational toolkit *N. oerstedii* relies upon to efficiently navigate back to its burrow, complementing its robust, yet error prone, path integration system with landmark guidance.

## Introduction

Stomatopods, better known as mantis shrimp, are benthic crustaceans renowned for their ballistic strikes and complex visual systems. As adults, most mantis shrimp species reside in shallow tropical marine waters, environments that are often structurally varied and therefore contain many potential visual landmarks [1]. In these environments, mantis shrimp typically occupy small holes or crevices for use as burrows, where they reside concealed for most of the day. During foraging, many stomatopod species leave the safety of their burrows for extended excursions, where they become vulnerable to predation [2-5]. Returning to the burrow efficiently is critical to minimize predation risk and to also reduce the chance that the vacated burrow will be claimed by another animal.

Mantis shrimp of the species *Neogonodactylus oerstedii* employ path integration to efficiently navigate back to their burrows between foraging bouts [5]. During path integration, an animal monitors the distances it travels in various directions from a reference point (usually home) using a biological compass and odometer. From this information, a home vector (the most direct path back to the reference point) is continuously calculated, allowing the animal to return to its original location [6-8]. As animals update their home vectors during excursions, small errors in odometric and orientation measurements are made. Over the course of an animal’s travel, these small errors accumulate in its path integrator. Therefore, with longer outward paths, increased errors of home vectors are expected [7, 9]. Path integration using idiothetic cues (those informed by stimuli anchored internal to the body) are particularly prone to accumulated error. As theory suggests, path integration in *N. oerstedii* is prone to this accumulated error [10]. To reduce this error, many path-integrators use landmarks to accurately pinpoint their goal [9, 11-14]. We hypothesized that in addition to path integration, *N. oerstedii* uses landmarks when available during navigation. The benthic habitats *N. oerstedii* occupy are structurally complex with an abundance of sponges, coral, rock, and seagrass to serve as potential visual landmarks (Fig. 1). Using landmarks during navigation would allow *N. oerstedii* to correct for error accumulated while path-integrating during foraging paths away from the burrow.

**Figure 1.**
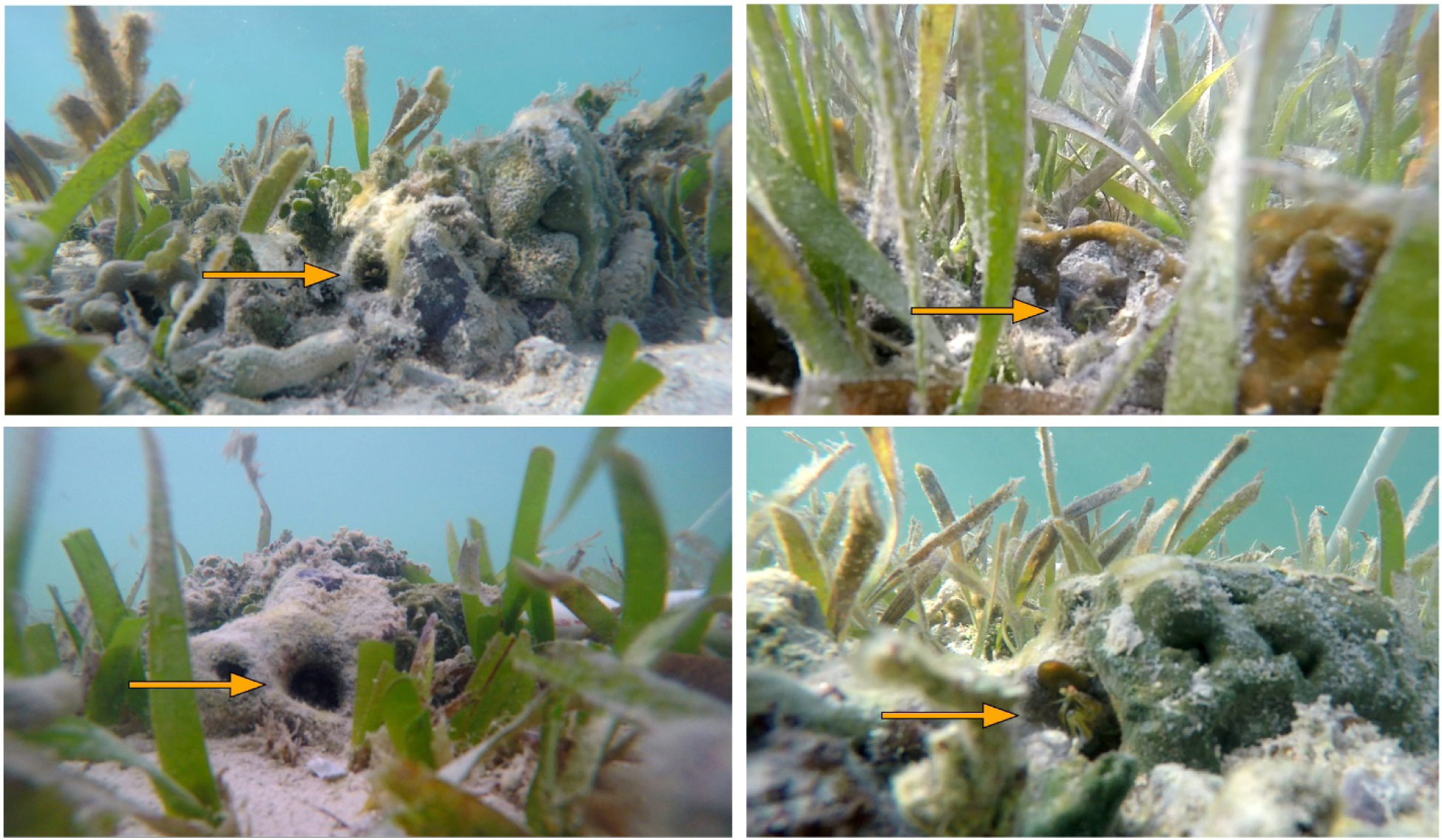
*Neogonodactylus oerstedii* inhabits shallow waters that offer an abundance of potential landmarks. Burrows are indicated by orange arrows. Note the abundance of potential landmarks, including marine vegetation, sponges, coral fragments, and rock rubble, available in the scenes. Stomatopods can be seen in their burrows in all except the bottom left panel, in which the photograph was taken when the animal had left its home.

## Results

### *Neogonodactylus oerstedii* uses landmarks during navigation

We placed *N. oerstedii* individuals in relatively featureless circular arenas filled with sand and sea water in a glass-roofed greenhouse. Vertical burrows were buried in the sand so that they were hidden from view when experimental animals were away. Snail shells stuffed with small pieces of shrimp were placed at one of two fixed locations approximately 70 cm from the location of the burrow in the arena (Fig. 2A). Foraging paths to and from the location of the food were video recorded from above.

**Figure 2.**
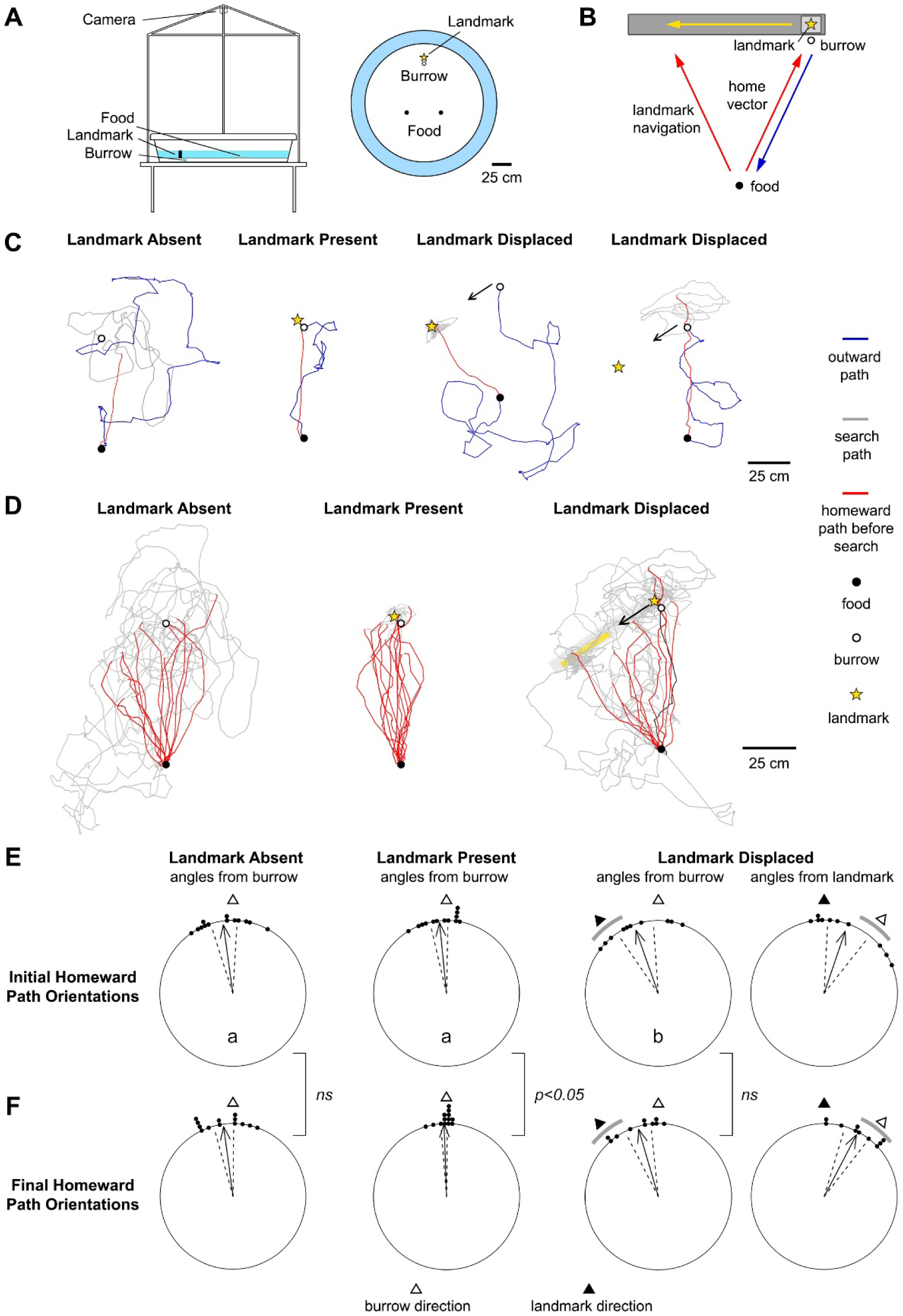
*Neogonodactylus oerstedii* uses a landmark to navigate back to its burrow while foraging. **(A)** Navigation arenas. Each arena was 150 cm diameter. A vertical burrow was set into the base of the arena 30 cm from the edge of the pool so it was invisible at range (empty circle). A landmark was placed adjacent to the burrow during some experiments (gold-filled star). Food was placed in one of two locations near the center of the pool (filled circles). Behaviors were video recorded from above. **(B)** Landmark displacement experimental design. Homeward paths were observed when a landmark adjacent to the burrow was displaced to a new location in the arena while experimental individuals were away foraging. **(C)** Examples of foraging paths from and to the burrow during the three experimental conditions. Blue lines represent outward paths from the burrow while red lines represent homeward paths before search behaviors were initiated. Grey lines represent homeward paths after search behaviors were initiated. Empty and filled circles represent the location of the burrow and food, respectively. Gold-filled stars represent the location of the landmark. Arrows represent paths of landmark displacements. **(D)** Data from all homeward paths. Lines and filled circles represent the same as in (C). The grey rectangle represents the track along which the landmark was displaced. The gold rectangle marks the range of locations to which the landmark was displaced during landmark displacement trials. The black tracing in the “landmark displaced” group marks the homeward path of an individual on its second run which, after orienting its initial homeward path towards the displaced landmark (in red), it returned to the food location and oriented towards the burrow (in black). **(E)** Orientations of homeward paths at one-third the beeline distance from the location of the food to the burrow (initial orientations). Each point along the circumference of the circular plot represents the orientation of the homeward path of one individual with respect to either the actual position of the burrow (empty triangle) or displaced landmark’s position (filled triangle). Grey arcs in the “Landmark Displaced” orientation plots represent the range of the directions of the either the displaced landmark or the burrow from at the location of the food. Arrows in each plot represent mean vectors, where arrow angles represent vector angles and arrow lengths represents the strength of orientation 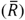. Dashed lines represent 95% confidence intervals. Different letters within orientation plots denote a significant difference between groups (p<0.05). “Landmark Absent” data were obtained from Patel and Cronin (2020a) [5]. **(F)** Homeward path orientations of groups same as in (E) measured immediately before search behaviors were initiated (final orientations).

As described by Patel and Cronin (2020a,b) [5,10], we observed that animals would make tortuous paths away from the burrow until they located the food placed in the arena. After animals located the food, they would usually execute a fairly direct home vector towards the burrow. If the burrow was not found using the home vector, animals would initiate a stereotyped search behavior (Fig. 2C and Extended Data Video 1).

To determine if *N. oerstedii* use landmarks during homeward navigation when available, a 2-cm diameter, 8-cm tall vertical cylinder with alternating 1-cm thick horizontal black and white stripes was placed adjacent to the burrow to serve as a landmark. Stripe cycles of the landmark would appear to span approximately 0.8 cycles/degree at the location of the food, approximately twice the visual resolving limit of *Gonodactylus chiragra* [15], a closely related mantis shrimp that can be slightly larger than *N. oerstedii*. Trials with the landmark present were compared to the results of previous experiments in which the landmark was absent [5].

Return trips in the presence of the landmark were more direct than trips in the landmark’s absence (P < 0.05; Fig. 2C-D and 3, and Extended Data Videos 1 and 2), supporting the hypothesis that *N. oerstedii* uses landmarks during navigation. This was primarily due to the virtual elimination of stereotyped search behaviors at the ends of homeward paths in the presence of the landmark. Instead, short directed searches for the burrow around the landmark were observed. Return trips were initially oriented similarly between the two groups (Groups were oriented: P < 0.001 for both groups; Orientations were not significantly different between groups: P > 0.5; All statistical outcomes are presented in Tables 1-3). However, during trials in the presence of the landmark, individuals appeared to correct for their initial homeward error over the course of the homeward path (P < 0.05), in contrast to what we observed in the absence of the landmark (P > 0.5; Fig. 2D-F). These results indicate that in the presence of a landmark, *N. oerstedii* uses both path integration and landmark navigation to navigate back to its burrow.

**Figure 3.**
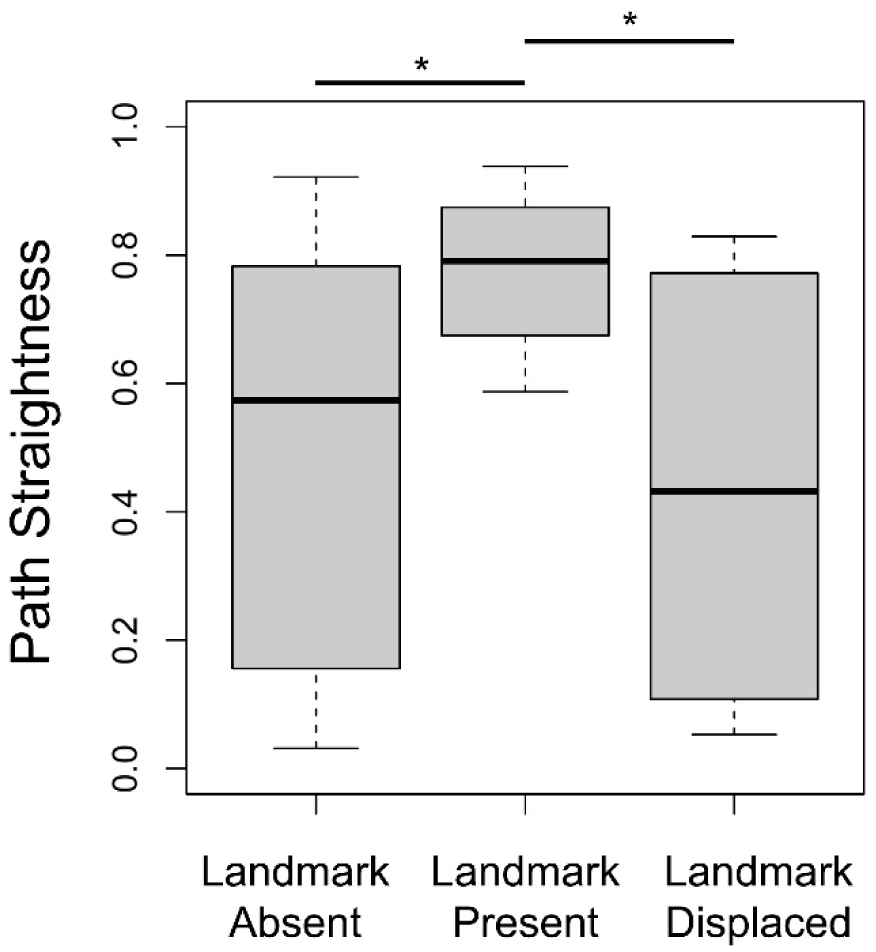
Homeward paths were more direct when a fixed landmark was present during navigation than when the landmark was absent or displaced to a new location in the arena during foraging. Straightness of homeward paths from the location of food to the burrow during trials when the landmark was present, absent, and displaced. Larger path straightness values indicate straighter paths with a value of one being a completely straight path from the food location to the burrow (a beeline path). Bars represent medians, boxes indicate lower and upper quartiles, and whiskers show sample minima and maxima. Asterisks indicate significant differences in path straightness between groups (P ≤ 0.05; Landmark Absent: n = 13, Landmark Present: n =13, Landmark Displaced: n = 10).

**Table 1:** Statistical outcomes of orientation analyses for all experimental groups. Orientations of homeward paths were measured relative to the burrow at one-third the beeline distance from the location of the food to the burrow (initial orientations) and were measured immediately before search behaviors were initiated (final orientations). Rayleigh tests of uniformity with Holm-Bonferroni multiple testing corrections were used to determine if groups were oriented. Data from this table can be viewed in Figure 2E and F.

| Experiment | P-value (uncorrected) | Holm-Bonferroni (corrected P-value) | n | $\bar{R}$ | Mean Vector Orientation ± S.E.M. |
| --- | --- | --- | --- | --- | --- |
| Landmark Absent (Initial) | <0.0001 | <0.001 | 13 | 0.949 | 354.4° ± 3.76° |
| Landmark Present (Initial) | <0.0001 | <0.001 | 13 | 0.974 | 352.2° ± 5.36° |
| Landmark Displaced (with respect to burrow position; Initial) | <0.0001 | <0.001 | 10 | 0.920 | 340.54° ± 7.76° |
| Landmark Displaced (with respect to landmark position; Initial) | <0.0001 | <0.001 | 10 | 0.894 | 18.79° ± 8.93° |
| Landmark Absent (Final) | <0.0001 | <0.001 | 13 | 0.966 | 352.32° ± 4.31° |
| Landmark Present (Final) | <0.0001 | <0.001 | 13 | 0.996 | 358.03° ± 1.47° |
| Landmark Displaced (with respect to burrow position; Final) | <0.0001 | <0.001 | 10 | 0.960 | 343.73° ± 5.44° |
| Landmark Displaced (with respect to landmark position; Final) | <0.0001 | <0.001 | 10 | 0.956 | 27.54° ± 5.74° |

**Table 2:** Summary of homogeneity of means circular statistical tests for orientation data. Comparisons of orientation groups in rows without an asterisk were analyzed using a Watson-Wheeler Test of Homogeneity of Means (test statistic is F). Comparisons of groups in rows with an asterisk (*) were analyzed using a non-parametric Watson’s Two-Sample Test of Homogeneity (test statistic is U^2^) since they did not adhere to the assumptions of a Watson-Wheeler Test. A P-value of less than 0.05 indicates a significant difference between groups. Data from this table can be viewed in Figure 2E and F.

| Experiment | P-value | Holm-Bonferroni | Test Statistic |
| --- | --- | --- | --- |
| Landmark Absent (Initial) vs Landmark Present (Initial) | 0.7355 | 0.7355 | 0.1168 |
| Landmark Present (Initial) vs Landmark Displaced (with respect to burrow; Initial)* | <0.02 | <0.04 | 0.2227 |
| Landmark Absent: Initial vs Final | 0.9827 | 1 | 0.000048 |
| Landmark Present: Initial vs Final* | <0.005 | <0.015 | 0.3373 |
| Landmark Displaced: Initial vs Final | 0.7414 | 0.7414 | 0.11234 |

**Table 3:** Summary of homogeneity of means statistical tests for path straightness data. The comparison in the row without an asterisk was analyzed using a paired T-test (test statistic is t). Since the “landmark displaced” group did not adhere to the requirements of a T-test, the row with an asterisk (*) was analyzed using a non-parametric paired Wilcoxon Signed-Rank Test (test statistic is V). The straightness of paths from groups within each comparison were significantly different from one another (P<0.05). The data from this table can be viewed in Figure 3.

| Experiment | P-value | Holm-Bonferroni | Test Statistic |
| --- | --- | --- | --- |
| Landmark Absent vs Landmark Present | 0.0216 | 0.0432 | 2.64 |
| Landmark Present vs Landmark Displaced* | 0.027 | 0.0432 | 49 |

### Mantis shrimp exhibit varied homeward paths when landmark navigation and path integration are placed in conflict

In light of the above results, we were interested in the confidence *N. oerstedii* places in its landmark navigation system when it is in conflict with its path integrator. In order to create this situation, homeward paths were observed when a landmark adjacent to the burrow was displaced to a new location in the arena while experimental individuals were away foraging. The landmark remained at roughly the same distance from the food location both before and after displacement. If *N. oerstedii* navigates using landmarks and trusts a landmark’s location over the location designated by its path integrator when homing, animals should orient towards the displaced landmark rather than the burrow’s location (Fig. 2B).

Homeward paths were less direct (P < 0.05; Fig. 3) and were differently oriented (P < 0.05; Fig. 2D-F) when landmarks were displaced compared to when they were left in place, further supporting the hypothesis that *N. oerstedii* navigate using landmarks. Some individuals oriented towards the displaced landmark while others ignored the displaced landmark, orienting towards the burrow (Fig. 2C and Extended Data Videos 3 and 4). Several individuals initially oriented towards the displaced landmark, but broke away from their initial trajectories during their homeward paths, orienting towards the burrow instead (Fig. 2D). Overall, however, differences observed between initial path orientations and the orientations of homeward paths at the end of the home vector were not statistically significant when the landmark was displaced (P = 0.36; Fig. 2E-F). One individual initially oriented its homeward path towards the landmark, only to turn around and return to the food location before adopting a revised homeward path oriented towards the burrow (Fig. 2D). These observations suggest that the path integrator of *N. oerstedii* is continually updated during foraging, even after homeward paths are initiated.

As just described, when landmarks were displaced some animals adopted paths initially oriented towards the displaced landmark while others ignored the displaced landmark completely, orienting towards the burrow. These results demonstrate that *N. oerstedii* must make decisions when the navigational strategies it relies on are in conflict and raise the question of how these decisions are made.

Due to errors inherit in path integration, *N. oerstedii* exhibit growing home vector errors with increased outward path lengths [10]. When the landmark was displaced, individuals may have evaluated this accumulated error during foraging, choosing to trust the position of the landmark when the accumulated error of the path integrator was high (i.e. confidence in the path integrator was low). However, we found that the orientations of homeward paths during landmark displacement experiments were not significantly correlated with the outward path lengths from the burrow to the food location (P = 0.16; Fig. 4A); nonetheless, the effect size of this relationship was fairly strong (r = -0.48), suggesting this hypothesis should not be completely discounted.

**Figure 4.**
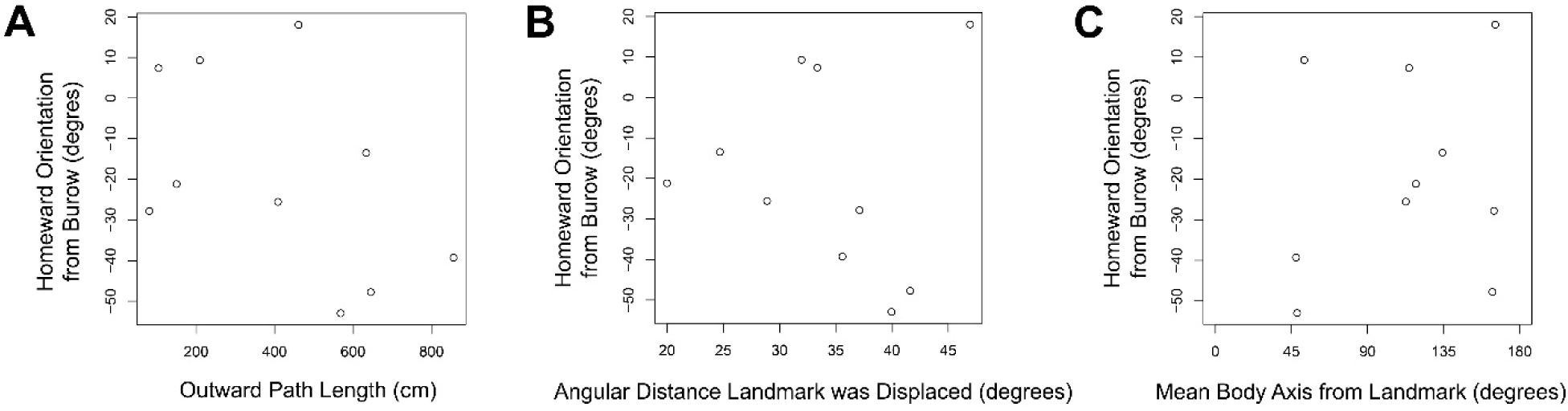
It is unclear why *N. oerstedii* chose to trust either the landmark or the home vector while navigating during landmark displacement experiments. **(A)** The orientations of homeward paths when the landmark was displaced was not significantly correlated with the length of outward paths from the burrow to the location of food (P = 0.16, n = 10, r = -0.48). **(B)** The orientations of homeward paths when the landmark was displaced was not correlated with the angular distance of landmark displacement along the track when viewed from the location of the food (P = 0.92, n = 10, r = -0.04). **(C)** Homeward path orientations were not correlated with body axis orientations of animals with respect to the landmark during its displacement (P = 0.604, n = 10, r = 0.19). Each point represents the mean body axis orientation of an individual with respect to the landmark measured at a sampling rate of 0.2 seconds during the landmark’s displacement.

Cataglyphid desert ants are model terrestrial species for studying navigation using path integration and visual landmarks. In experiments with these ants, when their path integrators are placed conflict with their surrounding landmark panorama, displaced desert ants will orient toward either the location indicated by their path integrator or toward a local landmark array depending on their distance from their nest, not on the error accumulated in their path integrators. These ants will orient using their home vectors, ignoring local landmarks, when displaced from at a distance greater than three meters from their nest; however, they will orient using the local landmark array when displaced from near the nest. When displaced from a distance of one meter from their nest, desert ants will orient with a mean vector not clearly directed at either their home vectors derived from path integration or the local landmark panorama, but somewhere in between [16]. Interestingly, orientation results of the desert ants displaced from roughly one meter from the nest are similar to those of *N. oerstedii* during the landmark displacement experiments described above. Stomatopods in those experiments were around 0.7 m from their burrows when initiating their homeward paths (Fig. 3E). These observations suggest that a cue integration mechanism resembling that employed by desert ants may also be present in mantis shrimp.

As an alternative hypothesis to account for the variation observed in homeward paths during experiments when the landmark was displaced, the deviation between the home vector and the landmark’s perceived position may have been at a preference threshold for either of the two navigation systems. For example, if the landmark was displaced further away from the burrow, the majority of animals may have trusted their home vector, while if the landmark was not moved as far from the burrow, the animals may have been more likely to trust the landmark’s position. However, when homeward path orientations during landmark displacement experiments were compared to the distance of landmark displacement along the track during those trials, no correlation was observed (P=0.92, r = -0.04; Fig. 4B). This suggests that the degree of landmark displacement did not influence the decision to orient toward the home vector or the displaced landmark during these trials.

Finally, we hypothesized that animals that may have observed the landmark’s displacement were more likely to disregard its location than those that may not have noticed displacement of the landmark. To investigate this hypothesis, we measured the orientations of all animals’ body axes with respect to the landmark while it was displaced, sampled at a rate of 0.2 seconds. We compared the means of these body axis orientations to the orientations of homeward paths and found no correlation (P = 0.604, r = 0.19; Fig. 4C). This suggests that either animals did not notice the landmark’s displacement or that observing the landmark’s displacement did not influence an animal’s decision to determine the burrow’s location by using the displaced landmark’s position or by using its home vector.

## Discussion

Our results demonstrate that *Neogonodactylus oerstedii* uses landmark navigation together with path integration while navigating back to its burrow while foraging. Landmarks are reliable references which can be used to correct for error accumulated by path integration; this is especially important during idiothetic path integration, which *N. oerstedii* uses when allothetic cues become unreliable [5].

Landmarks were used in a very basic situation during our experiments— as a beacon to home towards. Many other questions about how landmarks may be used by mantis shrimp arise from this work: Can mantis shrimp estimate the relative position of a goal to multiple landmarks? Do stomatopods use a snapshot mechanism like that employed by some insects to learn landmark arrays [13,17]? Do they possess cognitive maps akin to those thought to exist in mammals [18]? Do mantis shrimp learn to recognize landmarks encountered during foraging routes, exhibiting “trapline foraging”? Further, mantis shrimp are famed for possessing complex color vision, linear polarization vision in two spectral channels, and circular polarization vision [19]. Besides spatial vision alone, do stomatopods use these visual channels to identify landmarks? If so, how?

Mantis shrimp occupy a wide variety of marine habitats and depths, from structurally complex reefs to nearly featureless mud flats. Stomatopod species that occupy landmark-rich environments may weigh the importance of landmarks more heavily during navigation than stomatopods which occupy benthic environments relatively void of landmarks. Further, visual information rapidly attenuates with distance underwater due to extreme scattering of light in water. Therefore, the relative importance of landmark navigation over path integration may differ for mantis shrimp species occupying waters of different depths and turbidities.

Taken together with our previous work on mantis shrimp navigation [5, 10], this work offers an opportunity to study the neural basis of navigation, learning, memory, and decision making in stomatopods. Mushroom bodies, centers for arthropod learning and memory, are thought to play a prominent role in landmark learning in insects [20-23]. Prominent hemiellipsoid bodies, homologues of insect mushroom bodies, exist in stomatopod eyestalks [24]. As in insects, these neuropils may be crucial for navigation and landmark learning in mantis shrimp. A separate brain region, the central complex, plays a role in landmark orientation in *Drosophila melanogaster.* Here, landmark orientation is neurally based in the ellipsoid body of the central complex [25]. Stomatopods themselves possess a highly developed central complex composed of a collection of neuropils anatomically very similar to those found in insects [26]. Investigation of the function of stomatopod brain regions in light of our work may have implications for the evolutionary origins of navigational strategies and the neural architecture of the brain within the ancient Pancrustacean clade, a taxon which includes all insects and crustaceans [27], as well as in other arthropods.

In summary, *N. oerstedii* possesses a robust navigational toolkit on which it relies to efficiently navigate back to its burrow. First, *N. oerstedii* relies on path integration using multiple redundant compass cues to navigate back to its home [5]. If path integration does not lead *N. oerstedii* directly to its burrow, it relies on a stereotyped search behavior which is scaled to the amount of error it accumulates during its outbound foraging path to locate its nearby lost target [10]. Finally, the stomatopod will use landmarks, if available, to quickly pinpoint its target, offsetting error accumulated during path integration.

## Acknowledgements

We thank N.S. Roberts and J. Park for research assistance.

## Author Contributions

R.N.P. designed and conducted all research, analyzed all data, and prepared the manuscript. T.W.C. provided guidance and research support.

## Data and Materials Availability

The data that support the findings of this study are available from the corresponding author upon reasonable request. Correspondence and requests for materials should be addressed to R.N.P.

## Materials and Methods

### Animal Care

Individual *Neogonodactylus oerstedii* collected in the Florida Keys, USA were shipped to the University of Maryland, Baltimore County (UMBC). Animals were housed individually in 30 parts per thousand (ppt) sea water at room temperature under a 12:12 light:dark cycle. Animals were fed whiteleg shrimp, *Litopenaeus vannamei*, once per week. Data were collected from 13 individuals (5 male and 8 female). All individuals were between 30 and 50 mm long from the rostrum to the tip of the telson.

### Experimental Apparatuses

Four relatively featureless, circular navigation arenas were constructed from 1.5 m-diameter plastic wading pools that were filled with pool filter sand and artificial seawater (30 ppt; Fig. 2A). Arenas were placed in a glass-roofed greenhouse on the UMBC campus. The spectral transmittance of light through the greenhouse glass was nearly constant for all wavelengths, excluding the deep-UV-wavelength range (280 to 350 nm; Extended Data Fig. 1A). Celestial polarization information was transmitted through the glass roof of the greenhouse (Extended Data Fig. 1B-D). Vertical burrows created from 2 cm outer-diameter PVC pipes were buried in the sand 30 cm from the periphery of the arena so that they were hidden from view when experimental animals were foraging. Vertical 2 cm diameter, 8 cm high PVC columns with alternating 1 cm thick black and white horizontal stripes were placed adjacent to the burrows to function as removable landmarks. Stripe cycle widths of the landmarks were approximately twice the visual resolving limit of *Gonodactylus chiragra* (0.8 cycles/degree [13]), a closely related mantis shrimp that can be slightly larger than *N. oerstedii*, when viewed from the food location in the arena (a distance of 70 cm). Trials were recorded from above using C1 Security Cameras (Foscam Digital Technologies LLC) mounted to tripods placed above the arenas. During landmark displacement experiments, a thin 11 x 82 cm acrylic track with a movable platform was placed adjacent to the burrow (Fig. 2B). A landmark identical to the one used in trials in which the landmark was static, was mounted to the movable platform.

### Experimental Procedures

Individual *N. oerstedii* were placed in each arena and were allowed to familiarize themselves to the arena for 24 hours. During familiarization, the striped landmark was placed adjacent to the burrow, marking it during the animals’ initial explorations of the arena.

After familiarization, the landmark was either removed for trials in which the landmark was absent or left in place for trials in which the landmark was present. Empty *Margarites sp.* snail shells stuffed with pieces of food (whiteleg shrimp) were placed at one of two locations 50 cm from the periphery of the burrow. Each animal was allowed three successful foraging excursions (i.e. food placed in the arena was found) before foraging paths were used for analyses. If an individual did not successfully locate food within one week in the arena, it was replaced with a new individual.

During landmark displacement experiments, the landmark was carefully displaced along the track to a new location in the arena by the pulling of a thin fishing line tethered to the platform when animals were foraging away from their burrows. The distance from the food location to the landmark remained relatively constant while the landmark was displaced.

### Data and Statistical Analyses

Foraging paths to food locations and from them to the burrow were video recorded from above. In order to differentiate homeward paths from continued arena exploration, paths from the food locations were considered to be homeward paths when they did not deviate more than 90° from their initial trajectories for at least one-third of the beeline distance (the length of the straightest path) from the food location to the burrow. From these homeward paths, search behaviors were determined to be initiated when an animal turned more than 90° from its initial trajectory.

Paths were traced at a sampling interval of 0.2 seconds using the MTrackJ plugin [28] in ImageJ v1.49 (Broken Symmetry Software), from which the output is given as Cartesian coordinates. From these data, the inbound and outbound path lengths, beeline distances from food to burrow, and inbound and outbound indices of path straightness were calculated, where

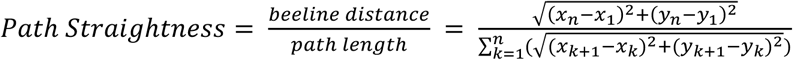

n = the last coordinate of the path

Additionally, the orientations of homeward paths when animals were both, at one-third of the beeline distance from the food source to the burrow (at which point the orientation of the home vector was usually observed) and at the end of the home vector (when search behaviors were initiated) were recorded using ImageJ.

We also measured the orientations of the body axes of all animals in respect to the landmark while it was displaced. These body axis orientations were sampled at a rate of 0.2 seconds. From these body axis orientations a mean body axis orientation was calculated for each individual.

Data from the “Landmark Absent” group in this study were taken from the “Not Manipulated” trials of the greenhouse experiments published in Patel and Cronin (2020a) [5].

All statistical analyses were run on R (v3.3.1, R Core Development Team 2016) with the “CircStats”, “circular”, “Hmisc”, and “boot” plugins. Orientation data were analyzed using the following procedures for circular statistics [29]. All reported mean values for orientation data are circular means. All circular 95% confidence intervals were calculated by bootstrapping with replacement over 1000 iterations.

As reported in Patel and Cronin (2020a) [5], no significant difference was observed between homeward orientations of males and females during experiments in the absence of a landmark (P > 0.5; Extended Data Fig. 2)), so data from both sexes were pooled for all experiments.

Rayleigh tests of uniformity were used to determine if homeward paths were oriented within a group for all trials. Parametric Watson-Williams tests for homogeneity of means were used to determine if those group orientations were significantly different from one another. The orientations of groups which did not fit the assumptions of the Watson-Williams test were instead compared using the non-parametric Watson’s two sample test of homogeneity. These tests were also used to compare differences between initial homeward path orientations (orientations at one-third the beeline distance from the food to the burrow) and final homeward path orientations (orientations at the initiation of search behaviors) for each group.

Homeward path lengths of trials in which the landmark was present were compared to those in which the landmark was absent using a paired T-test. A paired Wilcoxon signed-rank test was used to compare homeward path lengths of trials in which the landmark was static to those in which the landmark was displaced.

Pearson’s correlation tests were used for all correlative analyses.

Holm-Bonferroni multiple testing corrections were used for all tests when applicable.

## Extended Data Figures

**Extended Data Figure 1.**
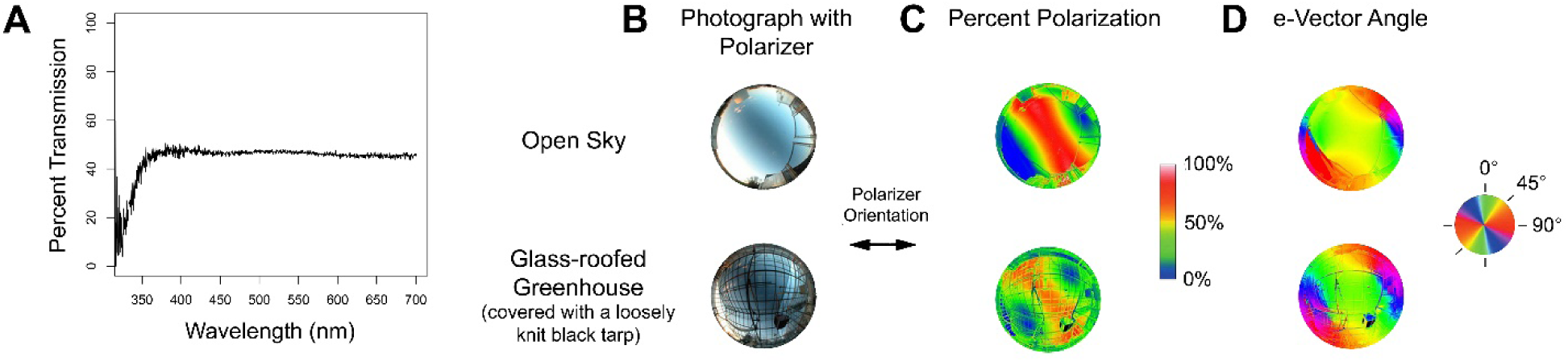
Photic conditions in the greenhouse where experiments were run. **(A)**Transmission of irradiance spectra through the glass-roof of the experimental greenhouse near sunset. The spectral transmittance of light through the glass roof of the greenhouse is nearly constant for all wavelengths greater than ∼360 nm. **(B-D)** Celestial polarization patterns are transmitted through the glass roof of the greenhouse**. (B)** Photographs of the sky at sunset on a day with very few clouds (November 24, 2015) using a fisheye lens and linear polarizer set in the east-west direction (as indicated by the arrow to the right of the photos). Photos were taken inside and outside the glass-roofed greenhouse used for experiments. **(C)** Percent polarization. Warmer regions in the images indicate higher percent polarization and cooler regions indicate lower percent polarization (see key). **(D)** e-Vector angle, indicated by the color corresponding the key to the right of the images. From Patel and Cronin (2020a) [5].

**Extended Data Figure 2.**
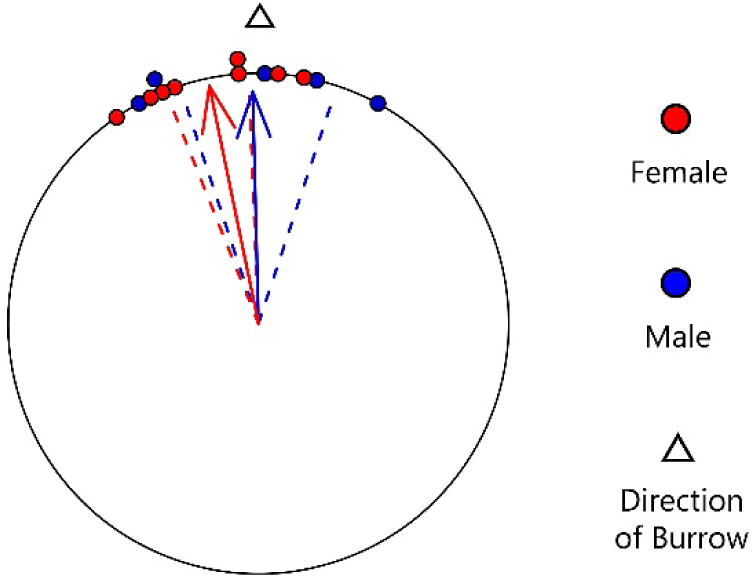
Male and female *N. oerstedii* orient towards home equally well while foraging. Homeward orientations of male and female individuals during experiments in the greenhouse when animals were not manipulated. Each point along the circumference of the circular plot represents the orientation of the homeward path of one individual with respect to position of the burrow (empty triangle). Blue-filled circles represent males while red-filled circles represent females. Arrows represent mean vectors, where angles of the arrows represent the mean vector angles and arrow lengths represent the strength of orientation in the mean direction 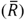. Dashed lines represent 95% confidence intervals. Males (n=5) and females (n=8) both exhibited significant orientations (p < 0.01 for both groups). No significant difference in orientation was observed between males and females (p>0.5). From Patel and Cronin (2020a) [5].

**Video 1. Foraging behavior of *Neogonodactylus oerstedii* showing homing in the absence of a landmark near the burrow.** Outward path is in blue, home vector path is in red, and search path is in grey. Filmed at 30 frames per second. Replay speed is indicated in the bottom-right corner of the video.

**Video 2. Foraging behavior of *Neogonodactylus oerstedii* showing homing in the presence of a landmark near the burrow.** Outward path is in blue and homeward path is in red. Filmed at 30 frames per second. Replay speed is in real time.

**Video 3. Foraging behavior of *Neogonodactylus oerstedii* showing homing after a landmark near the burrow had been displaced to a new location in the arena.** During this trial, the animal homed towards the displaced landmark. Outward path is in blue, home vector path is in red, and search path is in grey. Filmed at 30 frames per second. Replay speed is indicated in the bottom-right corner of the video.

**Video 4. Foraging behavior of *Neogonodactylus oerstedii* showing homing after a landmark near the burrow had been displaced to a new location in the arena.** During this trial, the animal homed towards its burrow. Outward path is in blue, home vector path is in red, and search path is in grey. Filmed at 30 frames per second. Replay speed is indicated in the bottom-right corner of the video.

